# LongReadSum: A fast and flexible quality control and signal summarization tool for long-read sequencing data

**DOI:** 10.1101/2024.08.05.606643

**Authors:** Jonathan Elliot Perdomo, Mian Umair Ahsan, Qian Liu, Li Fang, Kai Wang

## Abstract

While several well-established quality control (QC) tools are available for short reads sequencing data, there is a general paucity of computational tools that provide long read metrics in a fast and comprehensive manner across all major sequencing platforms (such as PacBio, Oxford Nanopore, Illumina Complete Long Read) and data formats (such as ONT POD5, FAST5, basecall summary files and PacBio unaligned BAM). Additionally, none of the current tools provide support for summarizing Oxford Nanopore basecall signal or comprehensive base modification (methylation) information from genomic data. Furthermore, nowadays a single PromethION flowcell on the Oxford Nanopore platform can generate terabytes of signal data, which cannot be handled by existing tools designed for small-scale flowcells. To address these challenges, here we present LongReadSum, a multi-threaded C++ tool which provides fast and comprehensive QC reports on all major aspects of sequencing data (such as read, base, base quality, alignment, and base modification metrics) and produce basecalling signal intensity information from the Oxford Nanopore platform. We demonstrate use cases to analyze cDNA sequencing, direct mRNA sequencing, reduced representation methylation sequencing (RRMS) through adaptive sequencing, as well as whole genome sequencing (WGS) data using diverse long-read platforms.

## Introduction

Recent advances in long-read sequencing technologies, particularly from Oxford Nanopore Technologies (ONT) and Pacific Biosciences (PacBio), allow users to sequence reads that are tens to thousands of kilobases (kb) long with high accuracy. Due to its momentous methodological advancement and broad application, long-read sequencing was chosen by the journal *Nature Methods* as “Method of the Year 2022” [1]. In 2023, Illumina also officially released the Complete Long Read technology, which yields high-accuracy long reads with an N50 read length of 5-7 kb [2]. These advancements make long read sequencing a viable option for a broad range of applications in genomics but require the development of novel bioinformatics tools which can quickly summarize important long read characteristics for quality control (QC) purposes. QC is a vital component of any sequencing pipeline to ensure that the raw data does not contain biases and technical artifacts, and to understand basic characteristics of the sequencing run. QC is of particular importance for long reads due to the relatively high base calling error rates [3], though the error rates have been dropping in recent years. In addition, these high throughput sequencing platforms require high-performance tools to quickly process large amounts of data: While short reads can produce reads ∼35-700 bases long [4], current long read sequencing reads are typically in excess of >10 kilobases[5]. A single ONT PromethION flowcell may generate up to 290 gigabases of sequence data, with up to 2.6 terabytes of signal data in FAST5 format [6]. Thus, long read QC tools must have the scalability to process the growing size of raw sequencing data in a fast manner to ensure that QC can be run as part of a routine sequencing pipeline.

There are several widely used and established short read QC tools that can be adapted for long read sequencing pipelines, such as FastQC [7], which provides an overview of raw sequence data as a summary HTML report and supports multiple data formats. However, FastQC does not provide mapping QC or important long read metrics such as N50 and read length distributions, and it has not been optimized to handle large datasets [7]. Thus, there is a need to develop a tool that enables fast, high throughput, and comprehensive QC summary statistics for important long read characteristics. In addition, due to a trade-off of increased error rates and lower throughput, it is often best to leverage a combination of sequencing technologies to achieve optimal results, which necessitates the need for a tool that provides QC metrics across platforms. Nevertheless, currently available QC tools for long-read data such as NanoPlot [8], PycoQC [9], NanoQ [10], NanoQC [8] and MinIONQC [11] typically support sequencing data formats for only one specific platform, such as PacBio or ONT, only support a subset of the major data formats, or focus on a specific aspect of the data, such as base modifications (Table 1) [8–12]. In addition, while available ONT QC tools all provide metrics on base called sequences, none currently provide support for summarizing signal intensity data or for extracting signals in specific genomic regions, which are important for epigenomics and epitranscriptomics studies.

**Table 1.** Comparison of filetype support vs. available long read QC tools (seqtxt = basecall summary files, uBAM=PacBio unaligned BAM).

| Tool | FASTA | FASTQ | BAM | uBAM | seqtxt | FAST5 | POD5 |
| --- | --- | --- | --- | --- | --- | --- | --- |
| LongReadSum | Y | Y | Y | Y | Y | Y | Y |
| NanoPlot | Y | Y | Y | Y | Y | N | N |
| PycoQC | N | N | Y | N | Y | Y | N |
| NanoQ | Y | Y | N | N | N | N | N |
| NanoQC | N | Y | N | N | N | N | N |
| MinIONQC | N | N | N | N | Y | N | N |

Here we present LongReadSum, a computational tool for fast, comprehensive, and high throughput long read QC: It supports data format types for all major sequencing technologies (FASTA, FASTQ, POD5, FAST5, basecall summary files, unaligned BAM and aligned BAM). To the best of our knowledge, it is also the first QC tool that can produce a summary for ONT basecall signal intensity data, and for the latest ONT POD5 format combined with BAM basecalling information.

## Implementation

LongReadSum is run from the command line as a non-interactive Python module and generates a comprehensive summary of different aspects of sequencing data in a timely manner by executing programs in a flexible multi-threaded C++ framework. The HTML report is generated in Python, while all QC metrics are computed in an extension module written in C++. The C++ module is wrapped for Python interfacing using SWIG. We leverage C++ multi-threading for BAM and basecall summary file analysis to achieve higher performance: On a 32-core computer with 4 Intel Xeon E5-4627 V2 3.3GHz 8 Core Processors with 1TB memory, QC metrics for an aligned BAM file (57 gigabases, with an N50 of 22 kilobases) from a single PromethION flow cell are completed in ∼15 minutes when running on 8 threads (Suppl. Fig. 1). For easy, platform-independent installation and use, we provide a Docker container option as well as an Anaconda package for Linux systems.

## Supported data formats

LongReadSum provides QC information for all major sequencing data formats, generating filetype-specific results as a summary text file and an HTML-based interactive report.

### FASTA and basecall summary files

FASTA is the simplest format containing a text-based representation of sequence records only [13]. ONT basecall summary files also contain basic sequencing read information. For data in these formats, the QC report contains basic statistics such as the total number of reads, base pairs, maximum, mean, and median read length, percent guanine-cytosine (GC) content, N50 (Fig. 1A), a read length histogram (Fig. 1B,C) and base counts for each nucleotide.

**Figure 1.**
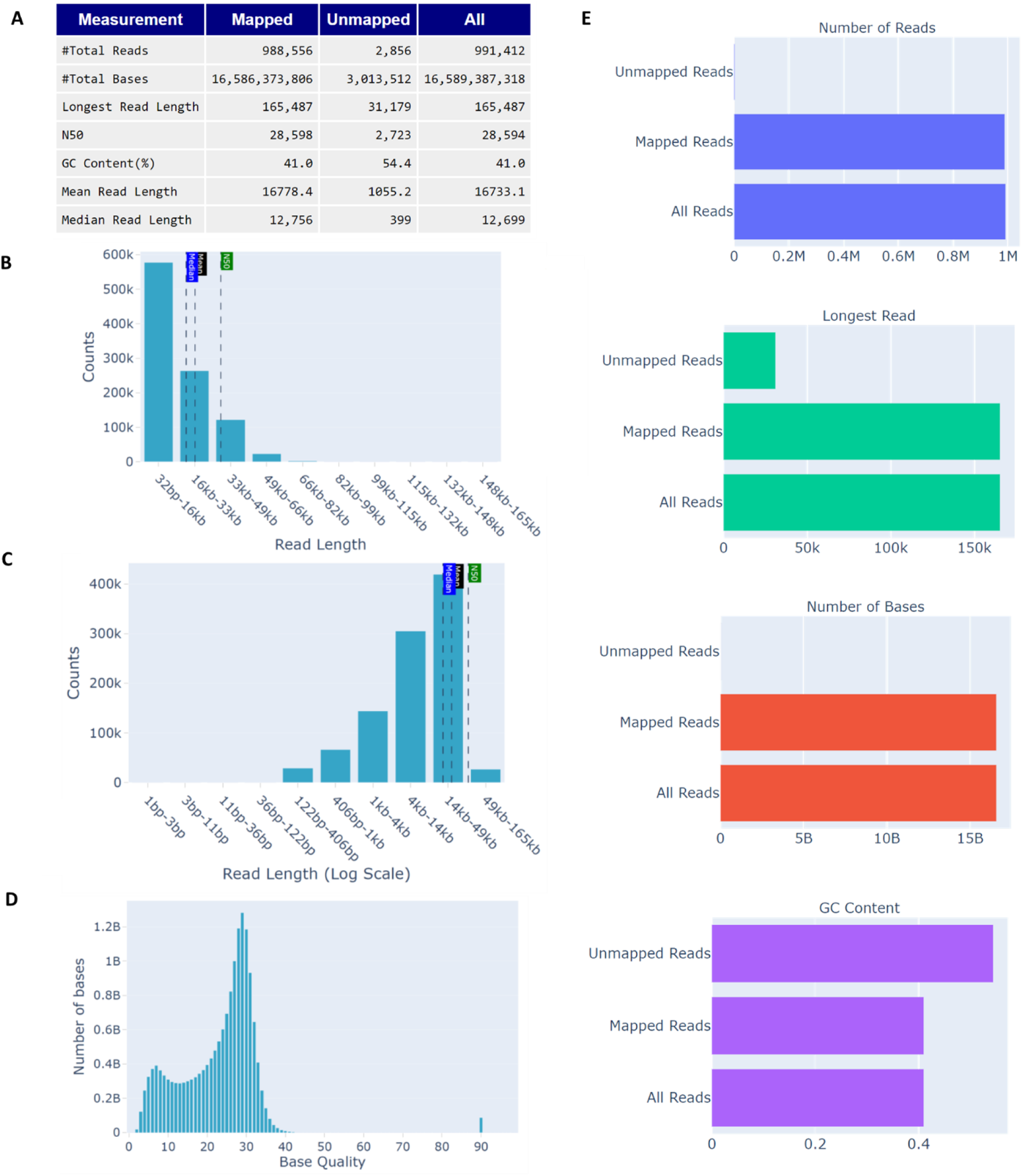
Basic QC metrics for a BAM file. (A) Table of read and base metrics; (B) Read length histogram; (C) Read length histogram (log scale); (D) Base quality histogram. The quality score of 90 is due to a potential bug in the Guppy basecaller from ONT; LongReadSum can optionally remove such erroneous calculations; (E) Basic statistics.

### FASTQ, FAST5, and BAM

The FASTQ, FAST5 and BAM (unaligned or aligned) formats builds upon FASTA to include sequencing read base quality information [14] where base quality is represented as PHRED scores [15, 16]. For these file formats with base quality information, the QC report generates a histogram of read average base quality, and overall base quality distributions (Fig. 1D).

### BAM with alignments

The BAM file format stores a binary representation of read alignments against a reference sequence [17]. For BAM files with alignment information, the report produces read and base alignment type counts, and mapped read and base statistics (Fig. 4). ONT BAM files may contain base modification information as well. While modkit[12] is an existing tool for summarizing ONT base modifications, LongReadSum provides a more comprehensive genome-wide summary if a FASTA reference genome file is provided.

### FAST5 with basecalls

ONT has its own proprietary FAST5 file type for recording time series signal data of pore ionic currents in addition to sequence reads and basecalls, and which follows the HDF5 file format [18]. There are currently available tools for FAST5 signal visualization and signal-to-base comparisons, which are useful for identifying signal anomalies in regions of interest [19–21]. Signal data has also proven useful at improving short tandem repeat (STR) detections [22–24]. In this tool, we include a FAST5 “signal” mode which generates interactive plots of raw pore signal intensity and corresponding base calls.

### POD5

The recently released ONT POD5 format is more efficient and is designed to replace the ONT FAST5 format [25]. It is different from FAST5 in that it does not contain sequence information, and thus we have designed LongReadSum to accept as input a POD5 and its corresponding basecalled BAM file, produced by e.g. ONT dorado [26]. This enables us to generate the signal-to-base correspondence as previously implemented in the FAST5 module.

## Results

### Performance

We compared LongReadSum performance with NanoPlot, the leading tool for comprehensive long read QC across major data formats. For testing, we used an aligned BAM file of a single PromethION flow cell from Oxford Nanopore’s open dataset of the HG002 genome (57 gigabases, with an N50 of 22 kilobases) [27]. We tested on a 32-core computer with 4 Intel Xeon E5-4627 V2 3.3GHz 8 Core Processors and 1TB memory. We ran both tools on the BAM file, and tested CPU and wall clock time performance using the GNU time program [28]. LongReadSum achieves shorter CPU processing and wall clock time by an order of magnitude: With 8 threads specified, QC results are generated in ∼15 minutes with ∼27 minutes of CPU processing time for LongReadSum, and in ∼1 hour 37 minutes of both CPU and wall clock time for NanoPlot (Suppl. Fig. 1).

### Basic Usage Case Report

Here we show an example BAM file QC output for a whole-genome sequencing (WGS) dataset of the HG002 genome obtained with the ONT MinION single molecule sequencing platform [29], base called with Guppy 6.1.2 and aligned to the GRCh38 reference. The HTML report includes a table of basic statistics for sequencing reads and bases including the N50, a useful statistical measure for describing read length and quality, as well as GC content, a useful indicator of DNA thermostability (Fig. 1A). In this sample, GC content is within the 40-60% range, and N50 is high relative to the mean and median read length. The report also includes interactive histograms for read lengths and base quality (Fig. 1B-D). Read length histograms help identify issues with sample preparation or sequencing which may result in reads of unexpected lengths (Fig. 1B, C). Expected read length distributions vary from ∼8-50 kb as expected based on the sample preparation [29]. A histogram of base quality across reads (Fig. 1D) as well as a histogram of average read quality score can be used to identify any biases in the preparation, whether the sequencing quality is consistent across reads, and any potential errors which may affect downstream analyses. In this sample, most base quality Phred scores are >20, indicating >99% accuracy for base calls. LongReadSum also yields metrics for both unmapped and mapped reads: The longest mapped read length and total numbers of mapped vs. unmapped bases as well as deviations in GC content distributions are important indicators of alignment accuracy and confidence (Fig. 1E).

### Use cases for different data types

Here we demonstrate different data type use cases, including analysis of cDNA sequencing, direct mRNA sequencing, ONT reduced representation methylation sequencing (RRMS), ONT whole-genome sequencing with base modification detection, as well as whole genome sequencing (WGS) in general. Both cDNA and direct mRNA involve the sequencing of smaller molecules that fall within several kilobases for long read sequencing with ONT, which can be verified in the read length comparisons for K562 cell line cDNA sequenced using MinION R9.4.1 flowcells (Fig. 2A) and K562 direct mRNA sequenced using PromethION R9.4.1 flowcells (Fig. 2B). ONT WGS library preparation has mean read length of ∼16kb (Fig. 2D, E). Publicly available Illumina Complete Long Read (ICLR) sequencing data of HG002 exhibits an expected N50 between 5-7 kb (Fig. 2F) [2]. LongReadSum also facilitates ONT RRMS studies, which involve sequencing within a targeted region of interest in a fraction of the genome. Our tool uses the resulting CSV file from RRMS to produce a separate QC report for accepted reads that pass the RRMS filtering criteria, as well as for rejected reads (Fig. 3). Accepted RRMS fragment sizes are around ∼5kb (Fig. 3C1), whereas rejected reads are typically less than 1kb as can be seen in the read length histogram (Fig. 3C2). The ONT R10.4.1 flow cell improves upon read accuracy when compared to R9.4.1. Although the R10.4.1 data was sequenced using Promethion, which produces over ∼6 times more sequencing data per flow cell than the MinION with the R9.4.1 flowcell [30], we can visualize the increase in sequencing read accuracy with R10.4.1 by comparing percentage of total reads with primary alignments (99.7% for R9.4.1 vs. 97.7% for R10.4.1), and percentage of total mapped bases with mismatches (5.63% for R9.4.1 vs. 4.14% for R10.4.1), insertions (1.46% for R9.4.1 vs. 1.15% for R10.4.1), and deletions (2.29% for R9.4.1 vs. 1.36% for R10.4.1) (Fig. 4). The source of these differences in error patterns can easily be visualized using IGV plots: R10 flowcells exhibit decreased mismatches and indels relative to R9 (Fig. 5). BAM files contain sequence alignment information, and LongReadSum provides a summary report of read and base alignment metrics, including a summary of each type of read and base alignment. High read and base alignment rates are indicative of high-quality sequencing data, and thus are important QC metrics.

**Figure 2.**
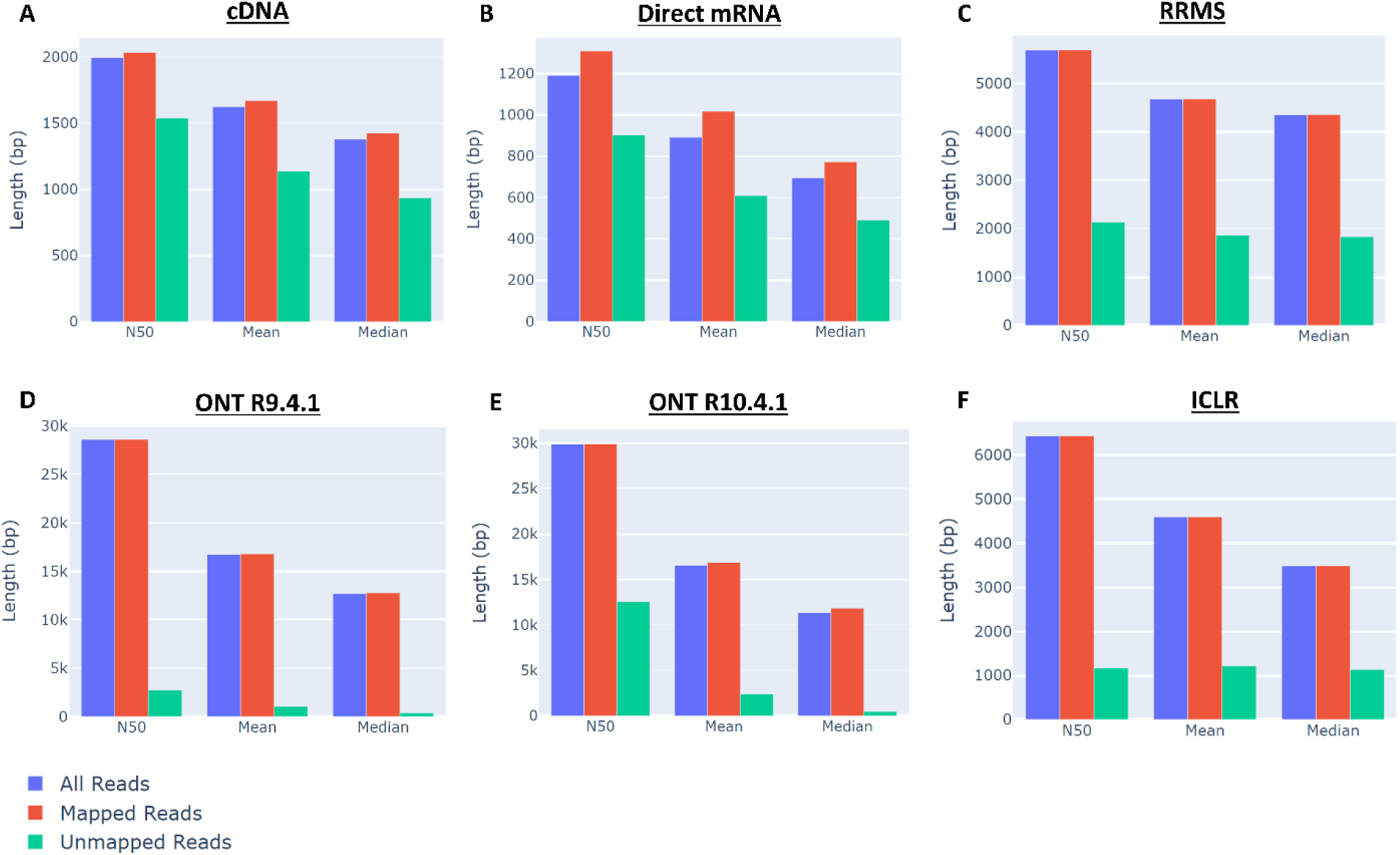
Comparison of read lengths for different types of data. (A) cDNA; (B) Direct mRNA; (C) Reduced representation methylation sequencing (RRMS). QC for accepted reads passing RRMS filtering steps are shown; (D) Whole-genome sequencing (WGS) with ONT R9.4.1 MinION flowcells; (E) WGS with ONT R10.4.1 PromethION flowcells; (F) WGS with Illumina Complete Long Read (ICLR) sequencing.

**Figure 3.**
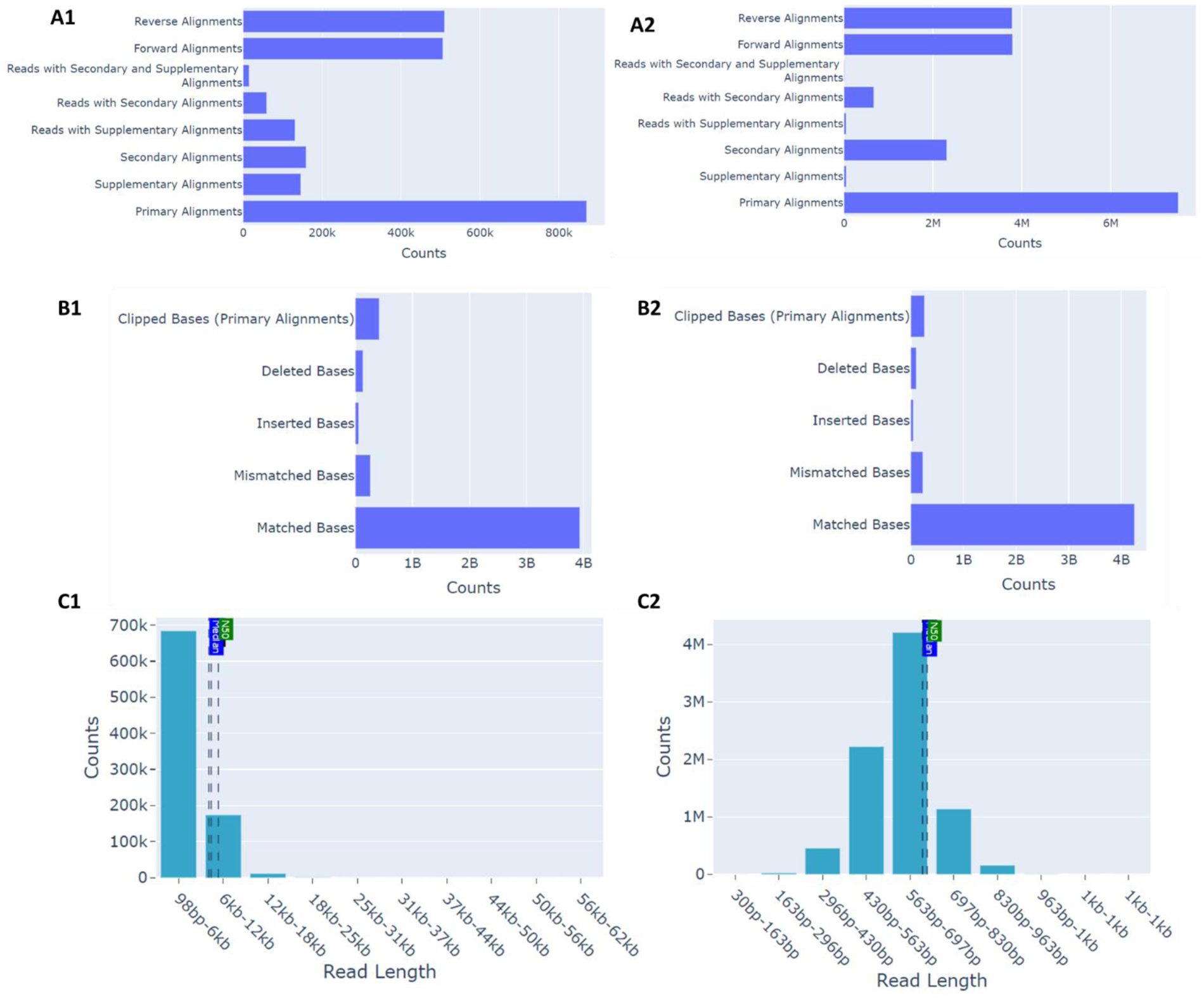
Comparison of RRMS accepted vs. rejected read statistics. (A) RRMS read alignment types for accepted (A1) vs. rejected (A2) reads; (A2) ONT R9.4.1 base alignment error rates; (B) RRMS base alignment error rates for accepted (B1) vs. rejected (B2) reads; (C) RRMS read length histogram for accepted (C1) vs. rejected (C2) reads.

**Figure 4.**
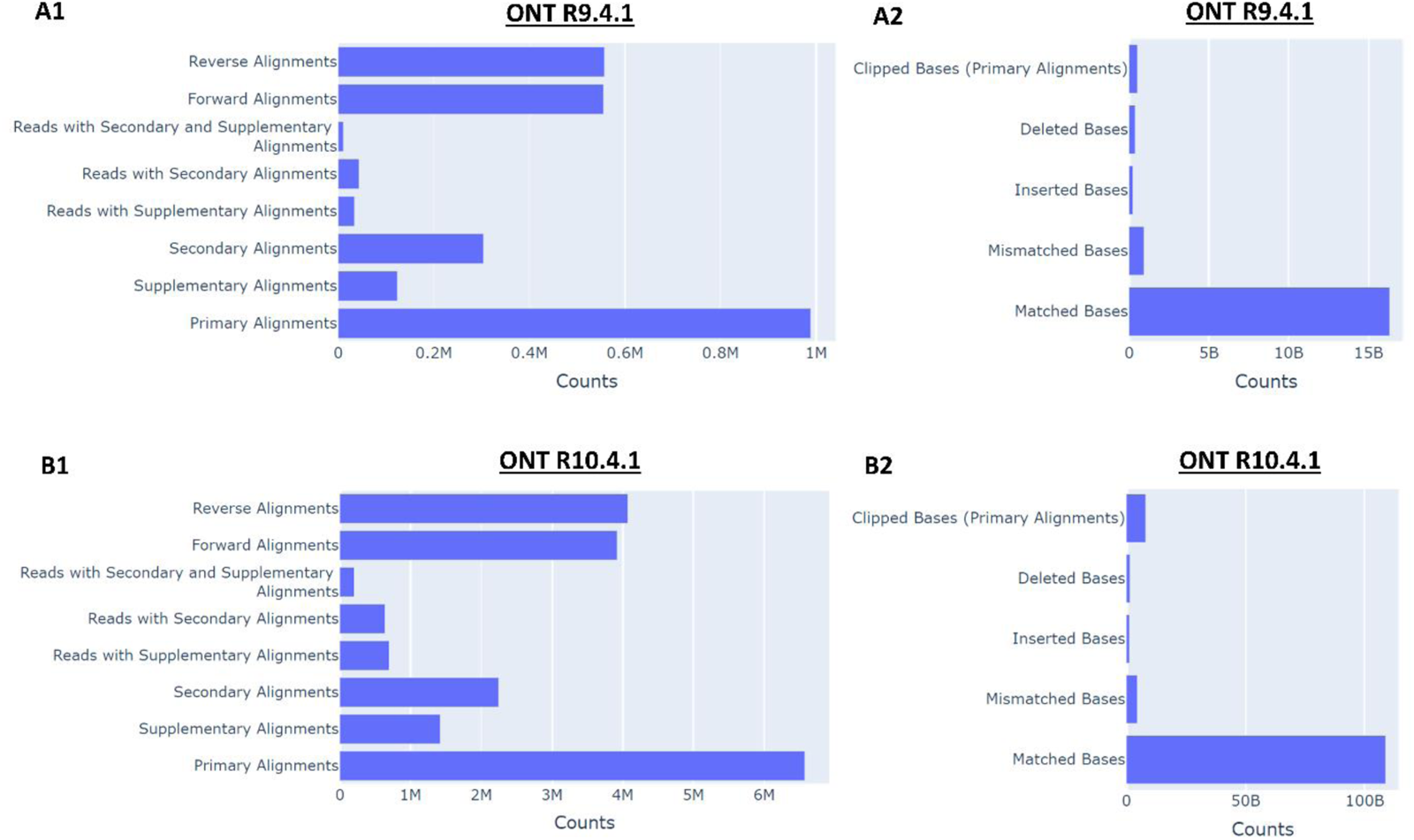
Comparison of alignment rates in whole-genome sequencing for ONT R9.4.1 MinION vs. R10.4.1 PromethION flowcells. (A1) ONT R9.4.1 read alignment types; (A2) ONT R9.4.1 base alignment error rates; (B1) ONT R10.4.1 read alignment types; (B2) ONT R10.4.1 base alignment error rates.

**Figure 5.**
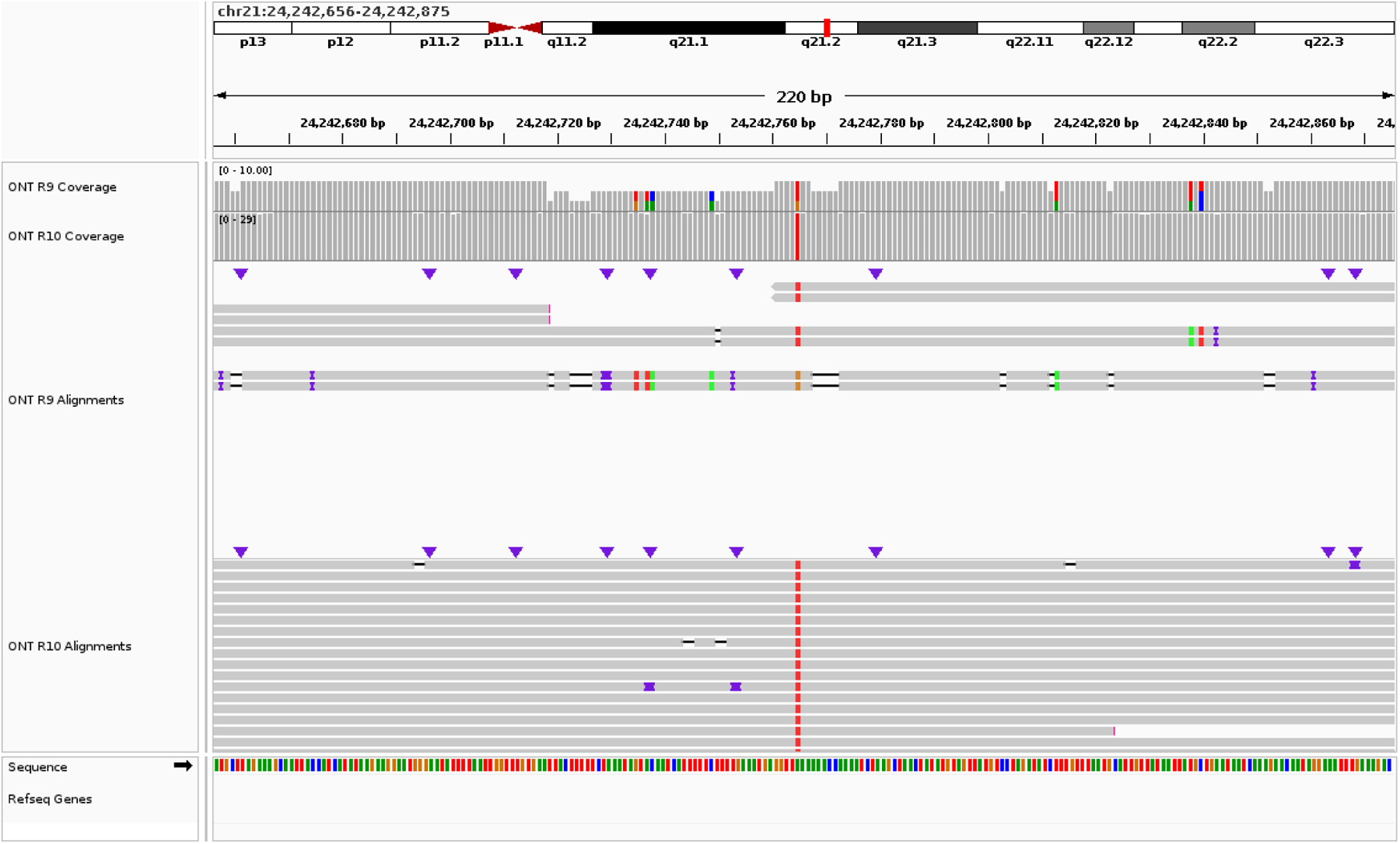
Comparison of error patterns in WGS data sequenced with ONT R9 vs. R10 flowcells visualized using IGV plots. Mismatches are indicated by colored bars. R10 flowcells exhibit increased read coverage and decreased s relative to R9

In Fig.6 we show how signal information from a FAST5 (or POD5) file can be used to validate the presence of regions of interest. In addition to storing read sequences, ONT FAST5 files store signal information generated during sequencing. These signals are used for base calling, and thus can be stored for future analysis with newer base calling algorithms. Thus, the signal data is highly valuable and can be used to identify patterns in regions of interest. For example, in Fig. 6 repeat motifs including [CAG]n (Fig. 6A), [GCC]n (Fig. 6B, 6C), as well as [C]n and [G]n (Fig. 6D) each exhibit stereotyped current signal patterns which can serve as evidence for validating true repeat region. Thus, a user can perform QC by visually examining read signal patterns to identify possible short tandem repeats or base call errors, for example by specifying a list of read names from a particular region in LongReadSum.

**Figure 6.**
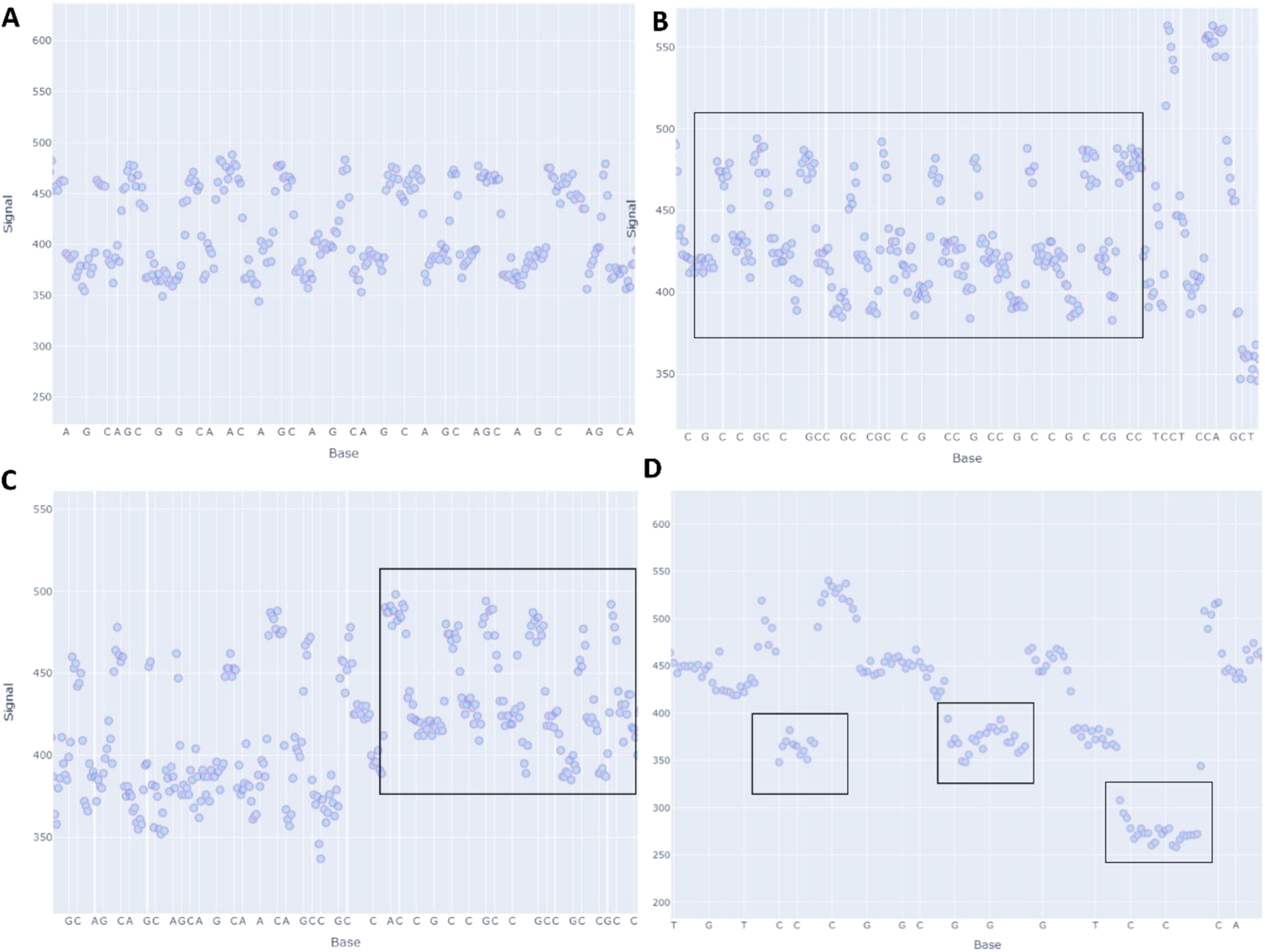
Repeat patterns in the FAST5 file ionic current signal data from ONT whole-genome sequencing are observable using LongReadSum. Specific repeat regions are indicated in the black box. (A) [CAG]n tandem repeats; (B,C) [GCC]n tandem repeats; (D) [C]3 and [G]3 repeats.

In Fig. 7 we show an example output for a publicly available POD5 file for HG002 sequenced with ONT R10.4.1 with its corresponding BAM file basecalled using dorado [26]. While the POD5 file contains the raw signal data, the BAM file contains the sequence and basecalling information that allows us to produce the signal-to-sequence correspondence plots for each read.

**Figure 7.**
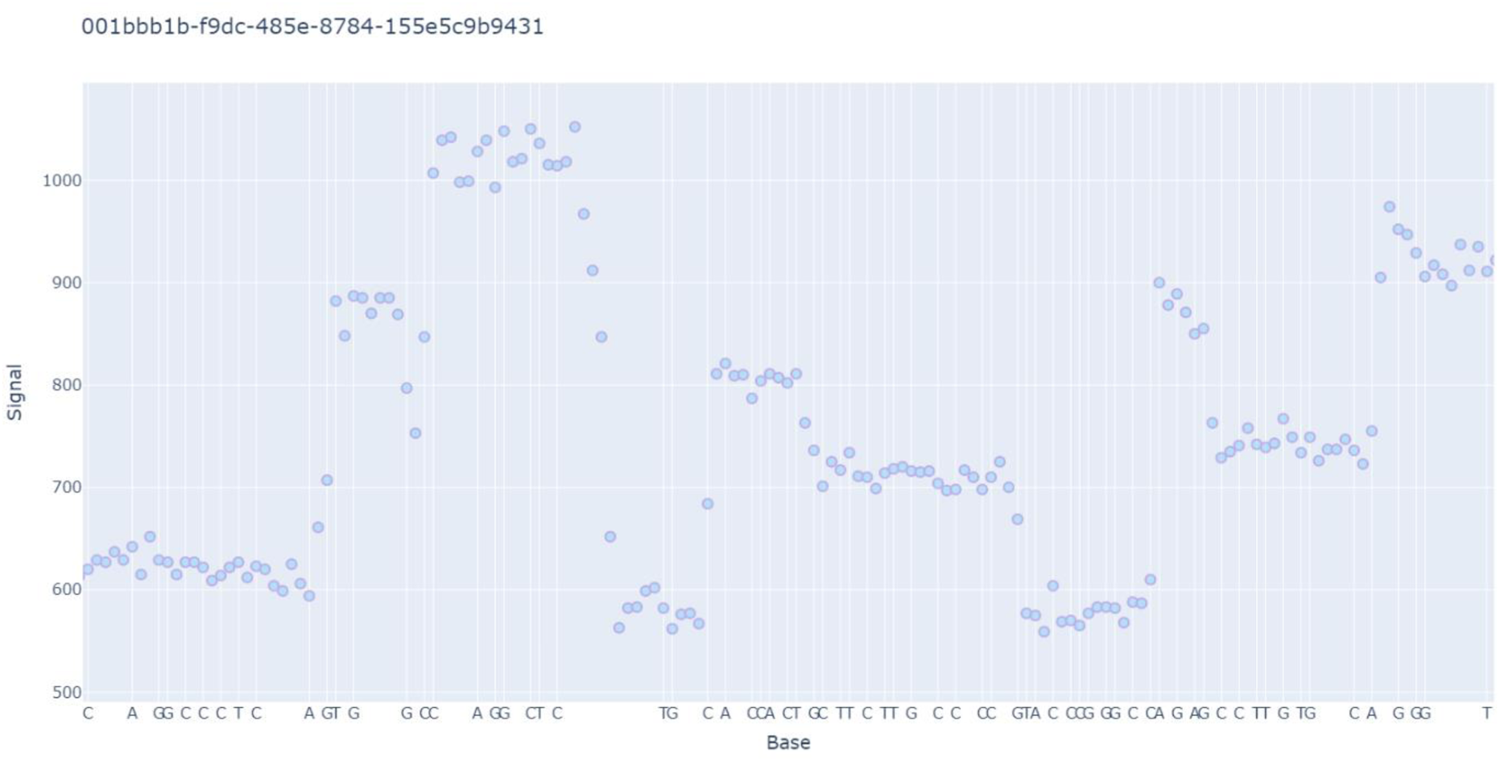
POD5 + basecalled BAM file ionic current signal data for a single read from ONT whole-genome sequencing are observable using LongReadSum. Data is HG002 sequenced with ONT R10.4.1.

In Fig. 8 we show an example LongReadSum base modification summary output for an ONT official public release of HG002 sequenced with MinION R9.4.1, basecalled with Guppy version 5.0.1, aligned to the GRCh38 reference genome, and with 5-methylcytosine (5mC) detection. We compare our results with modkit version 0.3.1, an ONT tool that provides similar metrics [12]. LongReadSum provides basic base modification information, including the total number of predictions, the total number of modified bases in the sample (combined, and for the forward and reverse strand) after filtering by a user-specified modification probability threshold, and finally the total modifications that align to CpG sites in the reference genome (Fig. 8). We compare results with modkit by running the following command, which yields CpG per-site modification values for a base modification threshold of 0.8:

**Figure 8.**
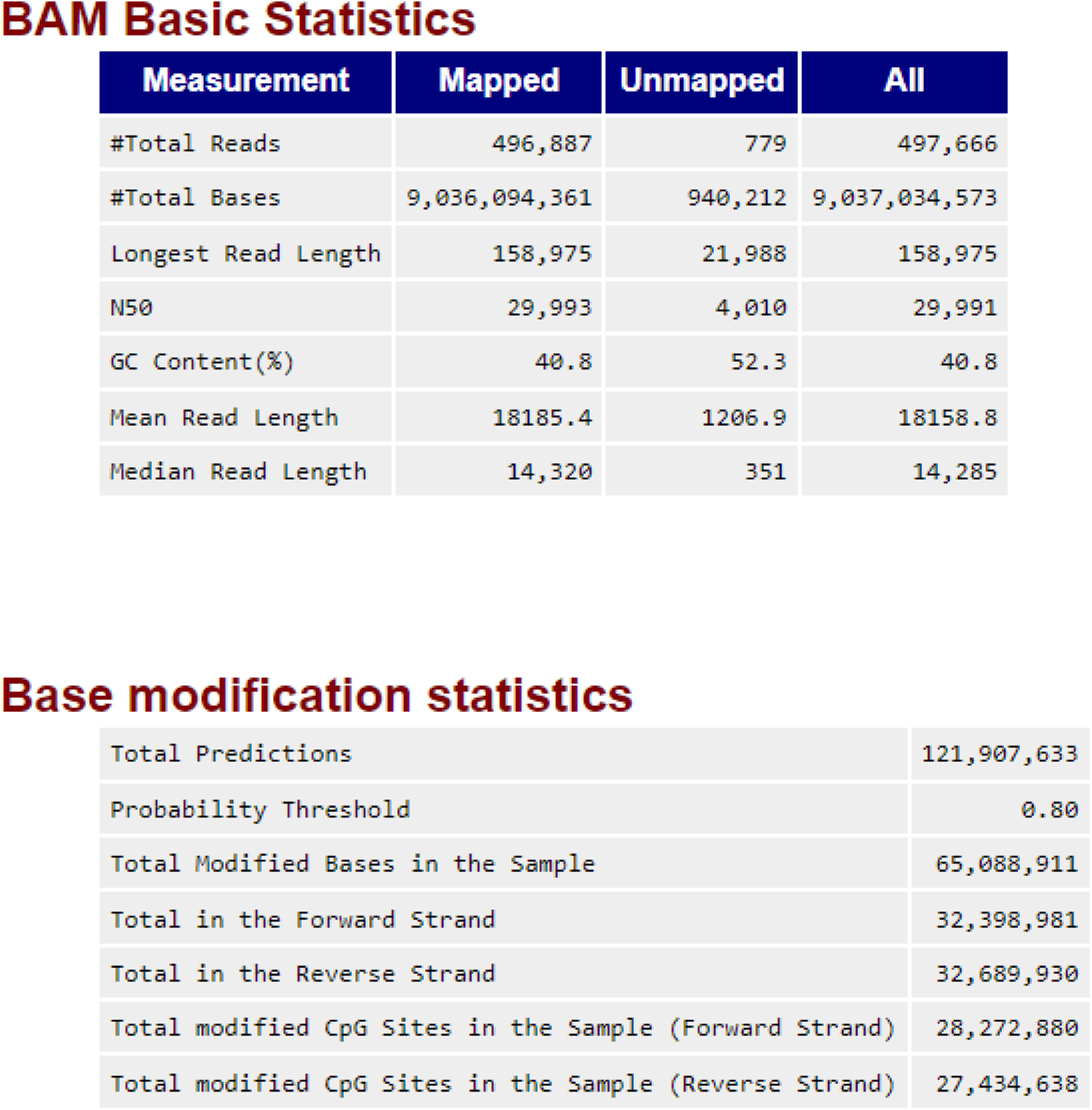
LongReadSum summary base modification and CpG site statistics from ONT whole-genome sequencing. Data is HG002 sequenced with ONT R9.4.1 MinION and 5mC base modification detection.

*modkit pileup $input_bam $output_txt --ref $ref_genome --filter-threshold 0.8 –cpg*

We then use a custom Python script to parse the modkit output file, which contains a list of unique reference positions with modifications, strand, and the number of reads supporting modification predictions (N_mod)_. Our resulting counts for modified CpG sites in the forward and reverse strand are identical to the modkit results obtained by summing over all values of N_mod_ for each strand.

## Discussion

LongReadSum is a fast and versatile tool providing a comprehensive overview of sequencing data and alignment quality, enabling the identification of significant errors that may preclude downstream analyses. It supports all major sequencing data formats, such as FASTA, FASTQ, FAST5, POD5, basecall summary files, unaligned BAM and aligned BAM files. It allows for quick identification of quality issues and biases in read distribution, base quality and mapping, and base modifications prior to downstream analyses. It also offers limited ability to summarize ionic current signals in specific genomic regions on the Oxford Nanopore platform.

### Future developments

In the future we aim to continue LongReadSum development to support evolving standards and file formats, as well as to provide support for emerging technologies often used in conjunction with long read sequencing, such as optical maps. BioNano optical maps have important applications in genomic analysis pipelines, including for *de novo* assemblies and structural variant calling. Thus, we will be adding QC support for optical map file formats including BNX and CMAP.

Finally, standards for sequencing file formats are continuously evolving with improvements in size, accuracy, and read/write speed to streamline downstream analyses. Although keeping up with evolving standards is a challenging task, we welcome these opportunities to expand our QC tool to incorporate support for new high-performance file formats which will benefit future users. In the future we also plan to include more in-depth mapping and base quality QC statistics for multiple sequencing runs on the same sample or for multiplexed samples in the same run. We hope that this tool is a useful addition to the community, in particular for researchers analyzing large sequencing datasets across multiple platforms. LongReadSum is released under an MIT license and is available at https://github.com/WGLab/LongReadSum.

## Acknowledgements

The authors thank all members of the lab for testing and evaluation of this tool. Research reported in this publication was supported by the NIH/NHGRI grant F31HG013259 and HG013359, and NIH/NIGMS grant GM132713. The content is solely the responsibility of the authors and does not necessarily represent the official views of the National Institutes of Health.

---

**Supplementary Figure 1.**
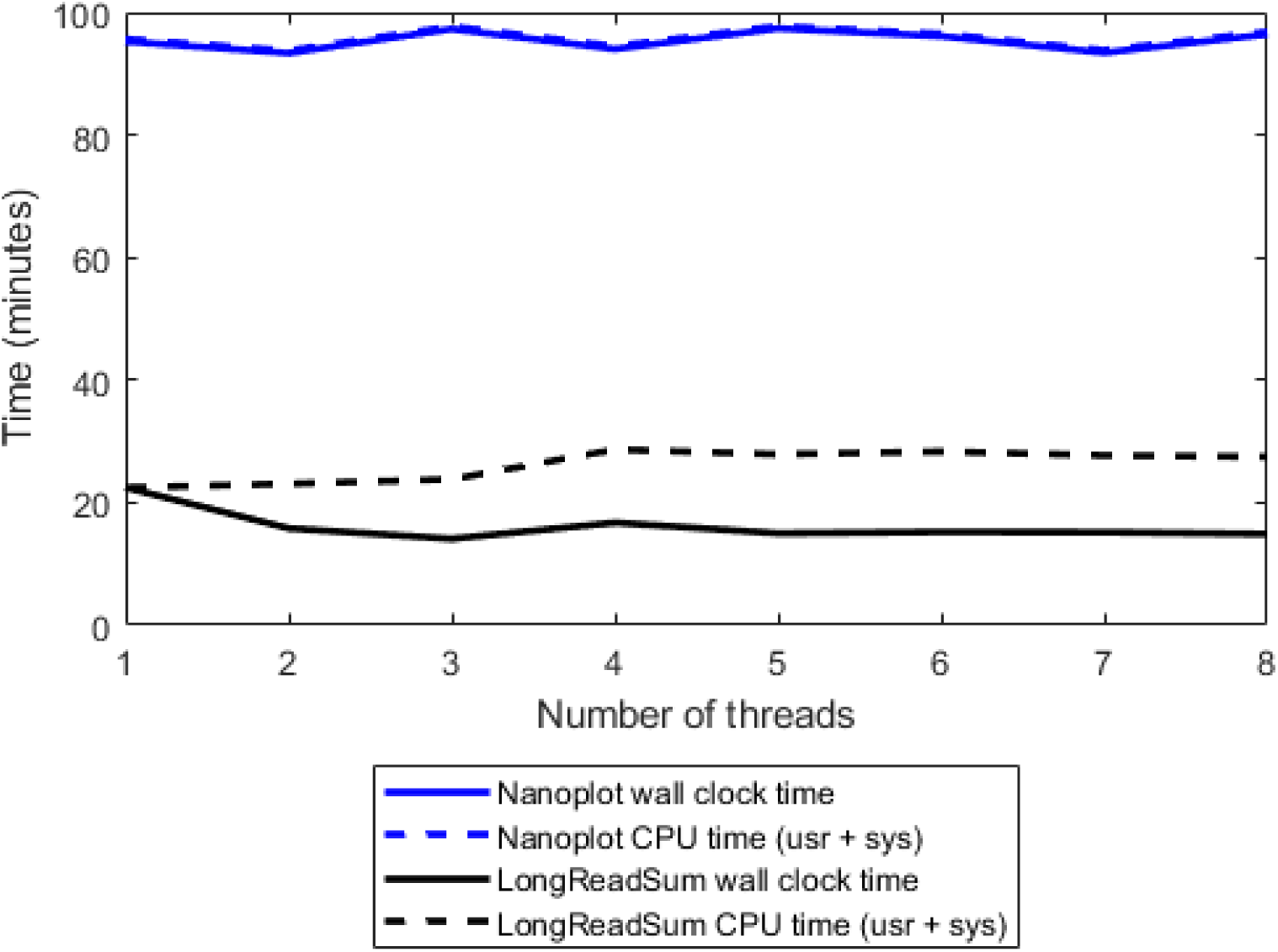
Comparison of multi-threading performance between LongReadSum and NanoPlot measuring time to completion of analyses (CPU vs. wall clock time). Data is an aligned BAM file from Oxford Nanopore’s open dataset of the HG002 sample sequenced with a PromethION R10.4 flowcell.

## References

1. 1. Method of the Year 2022: long-read sequencing. Nature Methods, 2023. 20(1): p. 1–1.

2. Illumina, I. Introducing Illumina Complete Long Read sequencing technology. 2023 [cited 2023 Aug. 17]; Available from: https://www.illumina.com/products/by-brand/complete-long-reads-portfolio.html.

3. Pollard, M.O., et al., Long reads: their purpose and place. Human Molecular Genetics, 2018. 27(R2): p. R234–R241.

4. Goodwin, S., J.D. McPherson, and W.R. McCombie, Coming of age: ten years of next-generation sequencing technologies. Nature Reviews Genetics, 2016. 17(6): p. 333–351.

5. Amarasinghe, S.L., et al., Opportunities and challenges in long-read sequencing data analysis. Genome Biology, 2020. 21(1): p. 30.

6. *PromethION 24/48 A100 IT requirements*. 2022.

7. FastQC. 2019; Available from: https://www.bioinformatics.babraham.ac.uk/projects/fastqc/.

8. De Coster, W., et al., NanoPack: visualizing and processing long-read sequencing data. Bioinformatics, 2018. 34(15): p. 2666–2669.

9. Leger, A. and T. Leonardi, pycoQC, interactive quality control for Oxford Nanopore Sequencing. Journal of Open Source Software, 2019. 4(34): p. 1236.

10. Steinig, E. and L. Coin, Nanoq: ultra-fast quality control for nanopore reads. Journal of Open Source Software, 2022. 7(69): p. 2991.

11. Lanfear, R., et al., MinIONQC: fast and simple quality control for MinION sequencing data. Bioinformatics, 2019. 35(3): p. 523–525.

12. *modkit.* Oxford Nanopore Technologies.

13. Pearson, W.R. and D.J. Lipman, Improved tools for biological sequence comparison. Proc Natl Acad Sci U S A, 1988. 85(8): p. 2444–8.

14. Cock, P.J., et al., The Sanger FASTQ file format for sequences with quality scores, and the Solexa/Illumina FASTQ variants. Nucleic Acids Res, 2010. 38(6): p. 1767–71.

15. Ewing, B. and P. Green, Base-calling of automated sequencer traces using phred. II. Error probabilities. Genome research, 1998. 8(3): p. 186–194.

16. Ewing, B., et al., Base-calling of automated sequencer traces usingPhred. I. Accuracy assessment. Genome research, 1998. 8(3): p. 175–185.

17. Li, H., et al., The Sequence Alignment/Map format and SAMtools. Bioinformatics, 2009. 25(16): p. 2078–2079.

18. The HDF5 Library & File Format. 2006; Available from: https://www.hdfgroup.org/solutions/hdf5/.

19. Loman, N.J. and A.R. Quinlan, Poretools: a toolkit for analyzing nanopore sequence data. Bioinformatics, 2014. 30(23): p. 3399–3401.

20. Ferguson, J.M. and M.A. Smith, SquiggleKit: a toolkit for manipulating nanopore signal data. Bioinformatics, 2019. 35(24): p. 5372–5373.

21. Payne, A., et al., BulkVis: a graphical viewer for Oxford nanopore bulk FAST5 files. Bioinformatics, 2019. 35(13): p. 2193–2198.

22. Fang, L., et al., DeepRepeat: direct quantification of short tandem repeats on signal data from nanopore sequencing. Genome Biology, 2022. 23(1): p. 108.

23. De Roeck, A., et al., NanoSatellite: accurate characterization of expanded tandem repeat length and sequence through whole genome long-read sequencing on PromethION. Genome Biology, 2019. 20(1): p. 239.

24. Giesselmann, P., et al., Analysis of short tandem repeat expansions and their methylation state with nanopore sequencing. Nature Biotechnology, 2019. 37(12): p. 1478–1481.

25. *POD5 File Format.* Oxford Nanopore Technologies.

26. *dorado*. Oxford Nanopore Technologies.

27. *Genome in a Bottle Ashkenazi Trio with Ligation Sequencing Kit V14*, O.N. Technologies, Editor.

28. Gordon, A., GNU Time. 1996.

29. Zook, J.M., et al., Extensive sequencing of seven human genomes to characterize benchmark reference materials. Scientific Data, 2016. 3(1): p. 160025.

30. *Advantages of nanopore sequencing.* Oxford Nanopore Technologies.

31. *modkit: Current limitations.* Oxford Nanopore Technologies.

